# Effects of mestranol on estrogen receptors expression in zebrafish

**DOI:** 10.1101/129502

**Authors:** Lianqiang Cheng, Xiaowen Wang, Shijun Li, Yongjun Liu, Zhengxin Guo

## Abstract

The estrogen receptor (ER) genes, which encode a group of important ligand-activated transcriptional factors, can modulate estrogen-target gene activities. Zebrafish (*Danio rerio*) have three ER receptor genes, *esr1*, *esr2a*, and *esr2b*. In this study, we examined the mRNA expression levels of these ER receptors after treatment with mestranol (EE3ME). Zebrafish larvae were exposed to 0.01, 0.1, 1, and 10 mg/L from 6 hours post-fertilization (hpf) and the mRNA expression levels of the ER genes were determined at 24, 48, 72 and 96 hpf. Treatment with mestranol led to a significant stimulation of *esr1* mRNA expression at lower concentration and reached maximum at 72 hpf, however, the esr1 mRNA levels were reduced at higher mestranol concentration during exposure. The gene expression of *esr2b* was markedly decreased and the *esr2a* remained unaffected at all concentration in the duration. Altogether, these results suggested mestranol might cause the disruption of endocrine activities in fish by mediating ER genes expression.

## Introduction

In all vertebrates, estrogens play important roles in many physiological processes, including general homeostasis and growth, and influence reproductive and nonreproductive target tissues [1-5]. These effects are mediated by numerous nuclear receptor proteins, the ERα and ERβ, which acting as ligand-activated transcription factors regulate estrogen downstream gene expression [6-9]. In vertebrates, three isoforms of ERs, ERα, ERβ and ERγ, have been cloned and characterized [10-12]. Estrogen activity is closely associated with endocrine disruptors and pharmaceutical estrogens in aquatic animals [13-16]. The disrupting chemicals cause severely endocrine dysfunctions in reproductive and developmental processes [17-21].

Mestranol, also knows as Ethynylestradiol 3-methyl ether, is a widely used synthetic steroidal estrogen [22]. Mestranol is the 3-methyl ether of ethynylestradiol, which is further converted to ethinylestradiol by demethylation in the liver [23]. The effects of estrogenic mestraol in the wildlife are largely unknown. Zebrafish (*Danio rerio*), a small tropical fish native to the rivers of South Asia and India [24, 25], has become one of the most important animal model in the research of developmental biology and toxicology [26-29]. Zebrafish have a wide variety of unique features including rapid development, easy maintenance, large number of offspring, transparency of embryos and access to experimental manipulation [30-32]. Many key genes and signaling pathways are highly conserved between zebrafish and advanced vertebrates [33, 34]. Due to many zebrafish lines of monogenic human genetic disease have been generated through forward genetic screens, supplying powerful tools to explore the basic cell biological processes that underlie the disease phenotype [35].

Zebrafish have three estrogen receptors, *esr1*, *esr2a* and *esr2b* [4, 36, 37]. The aim of this study was to determine the changes of estrogen receptors (ER) mRNA expression levels in zebrafish larvae treated with mestranol.

## Materials and methods

### Zebrafish husbandry

Zebrafish were raised and maintained as described [38, 39]. Zebrafish embryos were obtained by pair-wise mating of adult fish.

### Preparation of mestranol solutions

Mestranol (Sigma-aldrich) was dissolved in dimethyl sulfoxide (DMSO) (Sigma-aldrich) to make a stock solution (10 mg/mL). For the work solutions, to dilute 0.001, 0.01, 0.1 and 1 mL of stock solution into 1L aquarium water to make a final concentration of 0.01, 0.1, 1 and 10 mg/L, respectively. Zebrafish embryos were treated with different concentration of mestranol starting at 6 hpf, and samples were collected at 24, 48, 72 and 96 hpf. The control larvae were treated with same amount of DMSO in aquarium water.

### RNA extraction and cDNA synthesis

Total RNA was extracted from 50 embryos using the Direct-zol RNA MiniPrepKit (Zymo) according to the manufacturer’s instruction. One microgram of RNA was digested with 10 U DNaseI (NEB) to remove DNA contaminations for 30 min at 37 °C. The RNA was purified using OneStep PCR Inhibitor Removal Kit (Zymo) and the first strand cDNA was synthesized using the RevertAid First Strand cDNA Synthesis Kit (Thermo Fisher Scientific) according to the manufacturer’s protocol. The cDNA samples were diluted 10 times as template in the real-time PCR assays.

### Real-time PCR

Real-time PCR was performed as described somewhere else [40]. Briefly, qPCR mixture were prepared using iQ SYBR Green Supermix (Bio-Rad) and amplified using a CFX96 Touch Real-Time PCR Detection System (Bio-Rad). Primers specific for zebrafish esr1 (NM_152959.1), F-5’- GACTACGCCTCTGGATATCATTAC-3’ and R-5’- TGGTCGCTGGACAAACATAG-3’; esr2a (NM_180966.2), F-5’- GTCCGAGGTCTCAAGAGATAAAG-3’ and R-5’- CTTCCATGATCCGGGAGATTAG-3’; esr2b (NM_174862.3), F-5’- CAGACAACAGAGCCCAGAAA-3’ and R-5’- TCCTCTCGAAGCAGACTAGAA-3’; and β-actin (NM_131031), F-5’-TGCCCCTCGTGCTGTTTT-3’ and R-5’-TCTGTCCCATGCCAACCAT-3’. Primer specificity was confirmed by standard curve analysis and the mRNA expression of the three estrogen receptors was normalized to *β- actin*. Each experiment was carried out at least three times in triplicates. Calculated p values were considered significant at <0.05.

### Statistical analysis

The data were analyzed using the Statistical Package for the Social Sciences (SPSS) and one-way ANOVA with Tukey’s multiple comparison test was used for comparing multiple groups.

## Results

### Effect of mestranol exposure on ER expression in zebrafish larvae

To investigate the possible role of mestranol in zebrafish larvae, we treated the wild type embryos from 6 hpf to different concentration of mestranol (0, 0.01, 0.1, 1 and 10 mg/L). The mRNA levels of *esr1* were increased during the developmental stages from 24 to 96 hpf (Fig. 1A). At low concentration of mestranol (0.01 and 0.1 mg/L), the embryos showed increasing *esr1* mRNA expression during the treatment, and reached the maximum at the 72 hpf. However, the *esr1* mRNA expression was induced at 24 hpf but consistently decreased during the exposure at the concentration of 1 mg/L. Higher concentration (10 mg/L) of mestranol inhibited the *esr1* mRNA expression compared to the non-treated controls (0 mg/L). The mRNA expression levels of *esr2a* were slightly increased at low concentration of mestranol exposure (0.01 – 1 mg/L) at the first 72 hpf (Fig. 1B). However, the esr2a mRNA level were unchanged at higher concentration of mestranol treatment (10 mg/L) in the duration. At lower concentration (0.01 and 0.1 mg/L) of mestranol, the mRNA levels of esr2b were strongly induced during the exposure (24-96 hpf). However, higher concentration (1 and 10 mg/L) of mestranol inhibited the *esr2b* mRNA expression (Fig. 1C).

**Figure 1.**
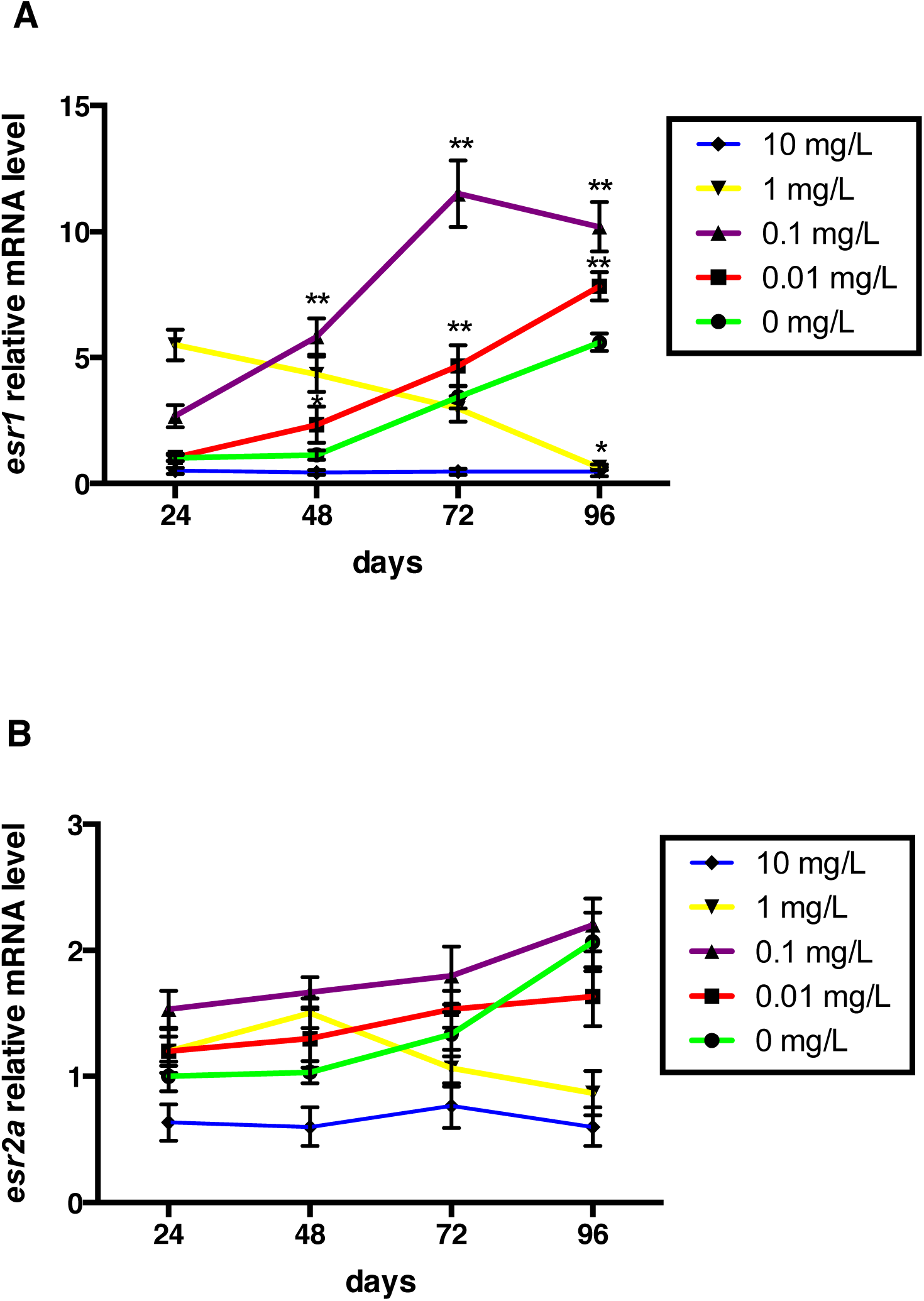
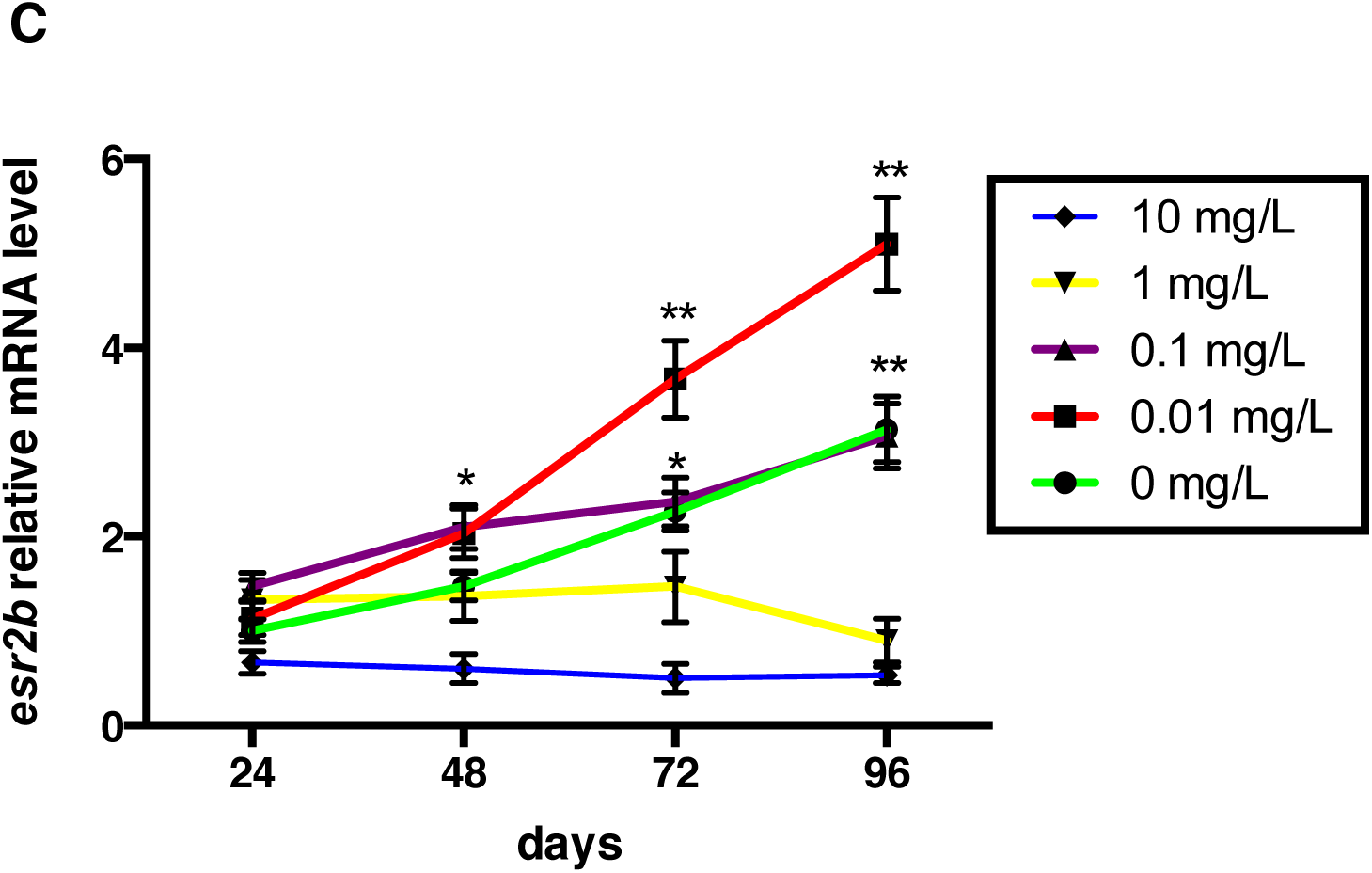
Effect of mestranol on *esr* expression levels in early-life stages. Expression levels of *esr1* (A), *esr2a* (B) and *esr2b* (C) genes in early larvae treated with different concentration of mestranol. Embryos were exposed to mestranol (0.01, 0.1, 1 and 10 mg/L) or non-treated control (0 mg/L) from 6 hpf and samples were collected at 24, 48, 72 and 96 hpf and processed for qPCR. Transcript levels after treatment were shown. Data were represented as mean ± SD of three independent experiments. * p< 0.05; **p<0.01.

## Discussion

This study investigated the in vivo effects of mestranol on the ER genes expression levels in zebrafish embryos. The expression levels of all three receptors in different tissues have previously been reported [41] and here we showed expression levels of these three genes in zebrafish larvae treated with mestranol. The mRNA expression levels indicated that the response in the zebrafish embryos of esr1, esr2a and esr2b was different when exposed to various concentrations of mestranol.

Many genes have been characterized that respond to ER in presence of ligand [42- 46], however, only a few number of genes have been identified in fish [12, 47]. The zebrafish has become a valuable animal model for investigating synthetic ligands that could affect the endocrine system.

The existence of two *esr2* genes may due to the gene duplication in zebrafish. These two genes have different expression levels possibly because of functional differences between *esr2a* and *esr2b*. The low *esr2a* gene expression levels in each group suggested that the *esr2* genes might have diverse functions during embryonic development. Treatment with lower concentration of mestranol induced expression of *esr1* and *esr2b* indicated that these genes were under regulatory control.

Environmental synthetic chemicals are generally considered as “weak estrogens” which could possibly be challenged by the potency they seem to have. Mestranol has the capacity to disrupt events during development and these effects are detected at lower concentrations that have no affecting survival. Zebrafish embryos are well suited as in vivo experimental model for evaluating molecular effects that exposed to mestranol.

